# Demographic Reporting and Phenotypic Exclusion in fNIRS

**DOI:** 10.1101/2022.11.08.515730

**Authors:** Jasmine Kwasa, Hannah M Peterson, Lietsel Jones, Kavon Karrobi, Termara Parker, Nia Nickerson, Sossena Wood

## Abstract

Functional near-infrared spectroscopy (fNIRS) promises to be a leading non-invasive neuroimaging method due to its portability and low cost. However, concerns are rising over its inclusivity of all skin tones and hair types (Parker and Ricard 2022, Webb et al 2022). Functional NIRS relies on direct contact of light-emitting optodes to the scalp, which can be blocked more by longer, darker, and especially curlier hair. Additionally, NIR light can be attenuated by melanin, which is accounted for in neither fNIRS hardware nor analysis methods. Recent work has shown that overlooking these considerations in other modalities like EEG leads to the disproportionate exclusion of individuals with these phenotypes – especially Black people – in both clinical and research literature (Bradford et al 2022, Choy 2020). In this article, we sought to determine if (1) biomedical optics developers and researchers report fNIRS performance variability between skin tones and hair textures, (2a) fNIRS neuroscience practitioners report phenotypic and demographic details in their articles, and thus, (2b) is a similar pattern of participant exclusion found in EEG also present in the fNIRS literature. We present a literature review of top Biomedical Optics and Human Neuroscience journals, showing that demographic and phenotypic reporting is unpopular in both fNIRS development and neuroscience applications. We conclude with a list of recommendations to the fNIRS community including examples of Black researchers addressing these issues head-on, inclusive best practices for fNIRS researchers, and recommendations to funding and regulatory bodies to achieve an inclusive neuroscience enterprise in fNIRS and beyond.

## 2. Introduction

Functional near-infrared spectroscopy (fNIRS) promises to be the leading non-invasive human neuroimaging method of the next few decades due to its portability, low cost, motion tolerance, and usability in special populations. This light-based modality was first ideated for blood-oxygenation estimation and has grown in its popularity, with publication counts doubling every 3.5 years (1,2). fNIRS is indispensable in many cognitive and psychological science settings, but especially in child development, hyperscanning, brain-computer interfacing, and other areas where movement and portability are challenges and which preclude EEG and fMRI as the leading non-invasive modalities (3–5).

As fNIRS increases in popularity, concerns over its inclusion of all skin tones and hair types are rising (6,7). While it has long been established that the physics of hair color, hair thickness, and skin pigmentation affect the detection of a NIRS signal (8), a systematic study is still missing that directly addresses the limitations of modern-day NIRS for different phenotypes. With these limitations, we are in danger of perpetuating bias against the darker skinned and thicker haired people of the world – individuals who already face racism and oppression worldwide. Here, we are careful to distinguish between phenotype and race: while phenotype refers to heritable physical characteristics such as hair and skin color, race is a social construct based on a collection of phenotypic, cultural, and regional indicators that hold power in society and affect the lived experiences of individuals who are minoritized and marginalized based on these indicators.

In this article, we briefly define technical limitations in biomedical optics for marginalized *phenotypes* and explore how they lead to disproportionate exclusion of people of marginalized *races* through a literature review. We sought to examine racial and phenotypic reporting specifically as compared to gender reporting, an established reporting category over the last few decades due to NIH mandated reporting. Although most guidelines combine “women and minorities”, we hypothesized that gender is reported at much higher rates than racial/ethnic demographics and treat it is a reporting exemplar.

## 3. Bias in fNIRS

### 3.1 Phenotypic Bias

fNIRS is used to measure real-time hemodynamics in the brain and is a proxy for brain activity. Red and near-infrared light is illuminated onto the scalp by a source optode and undergoes scattering and absorption throughout the underlying brain tissue until the attenuated light is detected at another optode some distance away from the source (see Figure 1). Two phenotypic “challenges” have emerged from this. The first is in *accessing the scalp* on individuals with coarse, dense, and curly hair; present-day optodes do not ensure that light sufficiently reaches the brain when thick hair occludes the scalp. The second challenge is in acquiring quality NIRS signals once the scalp is reached. Accurate measures of hemodynamics are impacted by the light absorption and scattering properties of the layers of tissue between the scalp and the brain, namely the dermis, skull, and blood vessels, and their particular tissue chromophores, including melanin (9,10). Variable melanin concentrations are not accounted for when calculating the hemodynamic response (11,12), potentially leading to inaccurate estimations of brain activity in darker individuals. Thus, phenotypic bias is perpetuated against darker skin, darker hair, and curlier hair.

**Figure 1.**
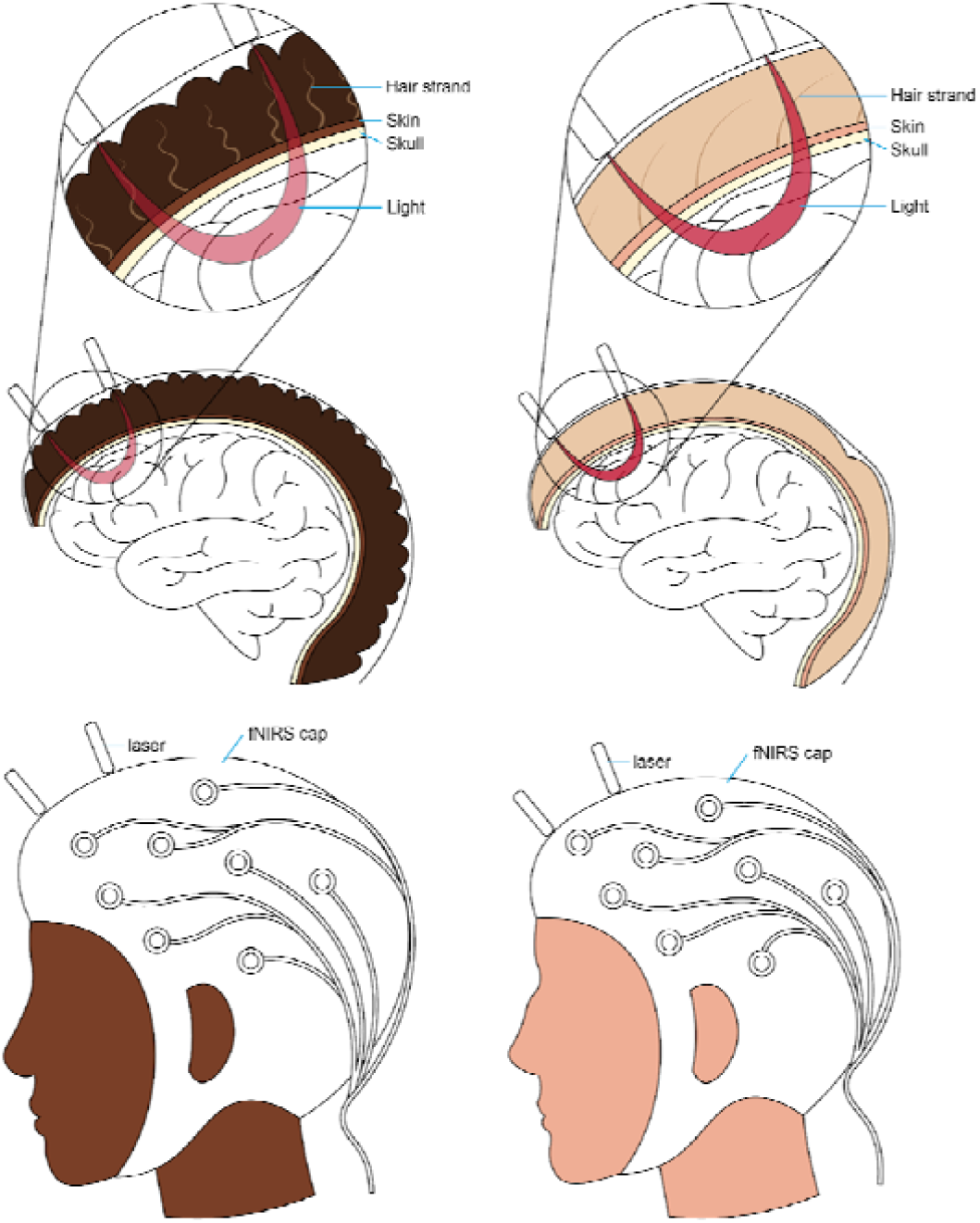
A combination of red and near-infrared (NIR) light at an optical source is shone into the brain non-invasively. From the source, light travels through the skin and into the brain surface before resurfacing at a detector or array of detectors elsewhere on the scalp. Using the scattering and absorption properties of NIR light in brain tissue, the relative amounts of oxygenated and deoxygenated hemoglobin present in the underlying brain region are calculated using the modified Beer-Lambert Law.

#### 3.1.1 Hair Type

One source of bias in fNIRS is its easier usability with short, straight, thin, and lighter-colored hair. Optodes must be as flush to the scalp’s surface as possible to get an optimal signal, and securely in place. Any optical obstruction between the fiber and the scalp, especially hair, can dramatically reduce the number of photons penetrating the scalp and ultimately the surface of the brain. Conventional NIRS systems cause concern for those with coarse, curly hair because the density and thickness of the hair may obstruct the fiber and because the caps may not accommodate the larger hair volume. Additionally, dark colored hair of any texture may introduce another source of melanin, which acts as a signal attenuator. Just as Etienne et al (13) found that traditional electrodes fail to maintain low impedance on individuals with coarse, curly, and dense hair, so too might fNIRS optodes fail to maintain physical contact with the scalp, since they are attached in the same fashion. Even with spring-loaded grommets and tension tops on fNIRS caps, anecdotally, the signal quality for participants with coarse and/or curly hair is poor. As a result, individuals with coarse, curly, and dark hair – often people of African, African-American, and Caribbean descent – are excluded from fNIRS studies (14–16). Therefore, fNIRS datasets tend to underrepresent Black and Brown individuals, which supports the need for our review. As a field we must ask: does the density, length, texture, or even the color of hair impact signal-to-noise ratio of the hemodynamics response inferred from fNIRS?

#### 3.1.2 Skin Pigmentation

Another source of bias in fNIRS is its better usability with lighter skin tones. Two key underlying assumptions in using the Beer Lambert Law are that hemoglobin is the main absorber in the dermis and that the tissue is optically homogeneous. In reality, several layers of the skin are optically heterogeneous, with melanin the dominating absorber of NIR light in the epidermis, and hemoglobin in the dermis. These oversimplifying assumptions lead to inaccurate estimations of chromophore concentration and an unreliable estimation of the hemodynamic response in individuals with skin pigmentation darker than a two on the Fitzpatrick scale, a spectrum of skin tones ranging from 1 (lightest) to 6 (darkest).

This inaccurate estimation of optically derived measures in different skin pigmentation levels is not new. The first clinically adopted NIRS device was the pulse oximeter, or pulse ox, used for non-invasive measurements of arterial oxygen saturation through the finger (17). Developed during WWII in the racially homogeneous Japan (18,19), the first pulse oximeter was adopted into clinical anesthesiology workflows in the 1980’s (17). While it has been long established that its accuracy is dependent on the calibration population (20), its design has not been reconsidered for darker skin. Recently, COVID-19 increased hospital and home-based pulse ox monitoring (21) leading to reporting that suggest skin tone may negatively affect accuracy (22,23). These limitations are currently under review by the FDA (24).

### 3.3 Exploring Exclusion

Methodological, experimental, and cultural limitations in current fNIRS practices contribute to what is called “convenience sampling” in brain imaging research. To accurately pinpoint convenience sampling in neuroscience research, we must assess the current phenotypic reporting practices in the theoretical and empirical neuroscience literature (5). In the following section, we present a literature review to determine current phenotypic and demographic reporting practices in fNIRS literature and conclude with a list of solutions to achieve an inclusive neuroscience enterprise.

## 4. LITERATURE REVIEW

### 4.1 Methods

In May and June 2022, we conducted a literature review of demographic and phenotypic reporting from articles in top English-language Biomedical Optics and Human Neuroscience journals. The three optics and two neuroscience journals were chosen to represent a range of articles covering fNIRS hardware and algorithm development and fNIRS as a tool in basic or clinical neuroscience research, respectively. Using PubMed, we saved a catalog of all articles in the given time range, selected journal name, and the keyword “fNIRS”. For the biomedical optics articles, we selected a 15-year time range; for the human neuroscience articles, we selected a 5-year time range. This time difference is because fNIRS’ adoption into basic research has understandably lagged fNIRS development; in all, both time ranges include the present day. Articles were retrieved on the open web or via subscription at the authors’ institution. For each article, we documented the number of participants, country of testing, any quantitative or qualitative reports of data exclusion, and participant demographics including: mention of sex or gender; mention of race, ethnicity, or nationality; mention of melanin, pigmentation, or Fitzpatrick scale; and mention of hair type. Animal and in silico studies were reported as “N/A”. We intentionally included sex/gender reporting in the analyses to compare as a baseline exemplar of “good” demographic reporting, since the NIH and many publishing bodies have encouraged or mandated reporting increasingly in the past 3 decades (25).

### 4.2 Results

From three top optics journals, we identified 110 articles from 2007 to 2022. We excluded *in silico* studies or those using animals, leaving a total of 90 articles with human volunteer participants (Figure 2A). While most studies reported gender (84.4%) as we predicted, nearly all articles failed to report phenotypic characteristics about participants being race/ethnicity (98.9%), skin pigmentation (96.7%), or hair type (93.3%). Over time, this trend does not seem to be improving (see Supplemental Figure 1).

**Figure 2.**
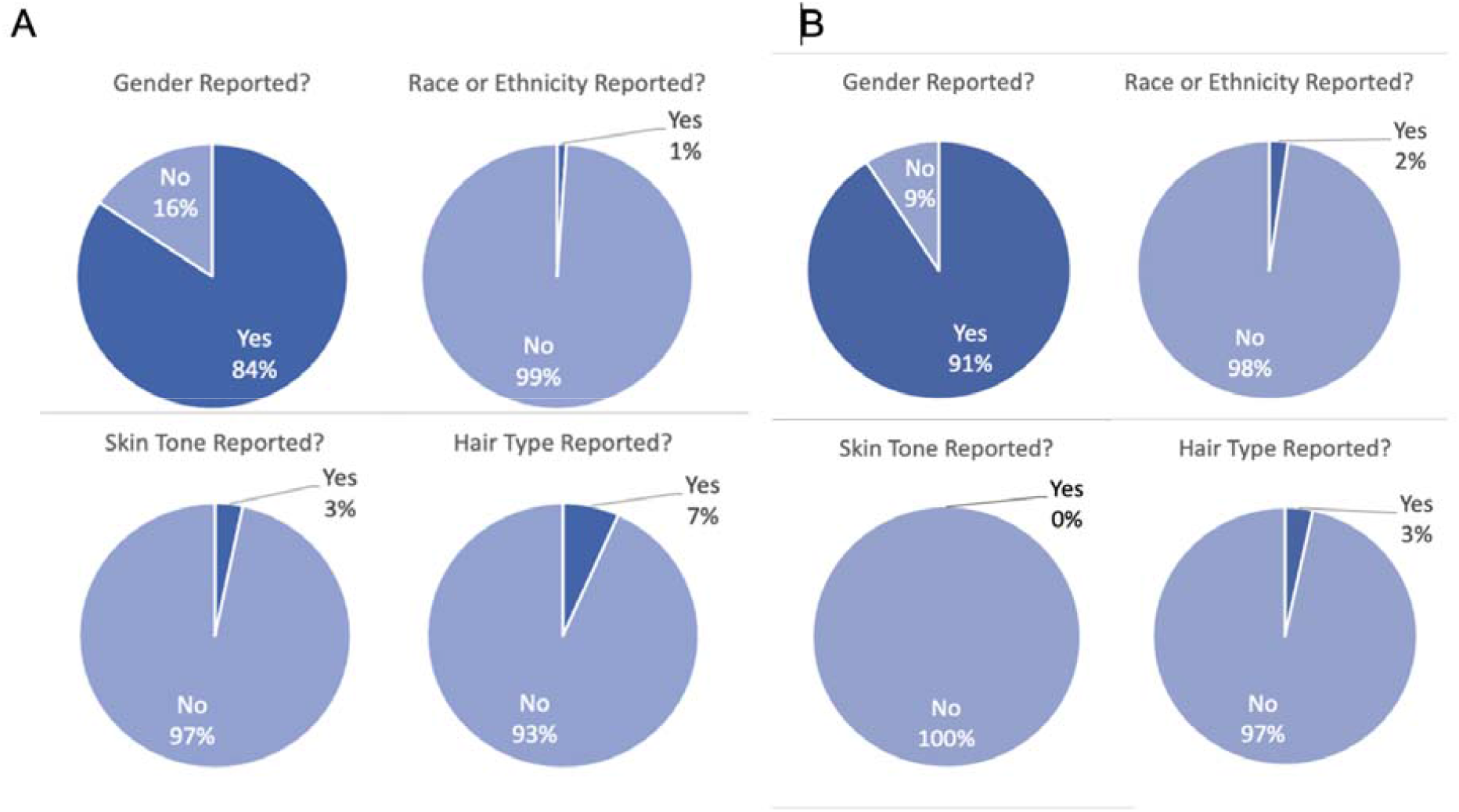
A) Demographic reporting for 90 articles with empirical human fNIRS data in three top biomedical optics journals. Overwhelmingly, gender is reported (“yes”) whereas race/ethnicity, skin pigmentation, and hair type are overwhelmingly not reported (“no”). B) Demographic reporting for 87 articles with empirical human fNIRS data in two top human neuroscience journals. Overwhelmingly, gender is reported (“yes”) whereas race/ethnicity, skin pigmentation, and hair type are overwhelmingly not reported (“no”).

We then repeated this analysis for two top human neuroscience journals that together publish a large proportion of basic science fNIRS articles. We identified 87 papers from 2017 to 2022 that used fNIRS as a tool (Figure 2B). Again, the vast majority of studies report gender (90.8%), but do not report race/ethnicity (97.7%), skin pigmentation (100%), or hair type (96.6%).

Lastly, we looked at the types of exclusion that were reported from all five journals. Only 69 of the 177 total articles (39.0%) mentioned if any participants were excluded *for any reason*. Of these 69, eight (11.6%) explicitly mention hair and four (5.80%) cite it as the main reason for the exclusion or withdrawal. The majority of the articles shared general reasons for dismissing a participant like “noisy data across channels,” “poor light shielding,” “technical issues… or low quality fNIRS data…,” “Bad fNIRS signal and technical issues,” and “poor cap fit.” The four articles cited thick or dark hair as being the reason for why a participant may have been excluded saying “poor data quality resulting from the subject’s relatively thick, black hair,” “unable to collect effective signals from fNIRS due to the participant’s thick, strong hair,” “had a lot of hair to obstruct light,” and “presumably due to dense and/or dark-colored hair.” No articles mention skin tone as being the primary source of signal noise. Race/ethnicity was the second least reported demographic and was typically reported by country of origin (e.g., “All participants were Chinese.”). A list of all the reasons for exclusion from the 69 articles are provided in the supplement (Supplemental Table 1).

Unfortunately, because of the low level of demographic reporting, we were not able to present data comparing the relative exclusion of marginalized and majority phenotypes.

### 4.3 Discussion

Our results point to two distinct issues: the under-reporting of exclusion and the potential, but unconfirmed, disproportionate exclusion of marginalized phenotypes. While recruiting diverse participants can prove challenging, simply reporting the participant makeup should be straightforward (see Section 5 for more discussion and recommendations).

In surveying biomedical optics journals, we sought to target work by the engineers who design fNIRS systems, those responsible for inclusive design practices. In surveying human neuroscience journals, we targeted work by end users. In both pursuits, we found that gender was reported in the vast majority of articles. This is likely due to the widespread adoption of gender reporting from NIH mandates that touched the animal research world as well as human research (and to our knowledge, other animals do not observe the social construct of race).

While four articles did report hair type and should be commended, there was only one article that explicitly mentioned hair type, hair color, *and* Fitzpatrick skin color. We especially commend that group for being transparent about the influences on their results and believe it should be the standard.

## 5. Recommendations

In the absence of adequate reporting of demographic data to determine exclusion trends in fNIRS literature, we think it’s important to highlight ways that the community – both developers and practitioners – can be more inclusive and more upfront about the inclusivity of their data. Based on our results, there is unequivocally exclusion based on, at minimum, the curliness and darkness of hair. To address this embedded bias, fNIRS tools and practices must change to accurately represent a heterogeneous population. The transition of fNIRS technology to more inclusive methodologies will require concerted efforts from engineers, scientists, clinicians, and imaging professionals following the example of groups already developing creative solutions.

### 5.1 Engineering Solutions

Some groups are actively addressing phenotypic bias limitations of fNIRS while maintaining other design requirements such as direct and prolonged contact with the scalp, maintenance of good signal-to-noise ratios, and increasingly higher spatial resolutions:

- A Texas-based group designed brush-type optodes to improve photon transmission and demonstrated its applicability with dark hair colors and high hair density by estimating power attenuation through a derived analytical model (26).
- More recently, Wood et al. began developing both novel inclusive optodes for curly hair *and* better algorithms to account for skin pigmentation. The team, which includes some authors from the aforementioned EEG work (13), has recently expanded to create novel pulse ox as well.
- A few studies mention personalized approaches to inclusive fNIRS setup, especially cap interfacing and design, a critical element to achieve quality optical contact. Sun et al. mounts light sources and detectors on a custom silicone cap to maintain contact (See supplemental figure 2) (27). The same group at the University of Michigan uses crochet hooks with LED lights to gently move hair during the optimization process before inserting optodes. While cap customization improves optode contact for different hair lengths and some hair types, the design may not be universal. For example, using crochet hooks can be painful for black hair as it tangles, and research assistants must be trained to do it.

### 5.2 Best Inclusive Practices for fNIRS Researchers

There are feasible approaches that researchers may consider to curb phenotypic exclusion and increase equity in the field.

#### Report demographics and phenotypes

We commend the one group in our sample that provided all demographic information upfront as well as the other groups that were honest about their exclusion of thick and coarse hair. Researchers involved in neuroimaging should explicitly report the racial and gender breakdown of their sample and, especially when there is exclusion of certain participants, describe the phenotypes such as hair color, hair type, and skin tone (28). Data about hair type and skin tone can be surveyed or judged by an experimenter familiar with the Fitzpatrick scales and hair typing scales such as the Andre Walker System or the L’Oreal system (14). Researchers should also consider the benefit of systematically quantifying the association of hair type, density, and melanin content of the scalp with fNIRS measurements. Formally defining these limitations through a systematic review will enable engineers to approach future advancements driven by these factors.

#### Adopt inclusive methodologies and hire a diverse research team

Although fNIRS systems need improvements, there are other reasons why darker skinned and curlier haired individuals are excluded from psychological research and design solutions. Many standard procedures foster an unpleasant environment and result in voluntary participant withdrawal from marginalized backgrounds especially for special populations in which fNIRS is beneficial. For example, children with darker pigmented skin and curlier hair textures (and their parents) may get frustrated and lose trust in the researchers because of the complex setup process, which involves repeatedly moving the cap and hair. Moreover, individuals with intellectual disabilities – a large proportion due to fNIRS’ portability and motion tolerance – may not be able to handle the inconvenience.

To improve participant experience, researchers should train to work with a range of hair types as standard practice. Adverse outcomes of unpreparedness include longer setup times, microaggressions, participant discomfort, and participant dropout. fNIRS researchers should consider developing guidelines for preparation that will serve as standard operating procedure. Given some similarities in configuration and setup between EEG systems and fNIRS equipment, following Etienne et al.’s suggestion for adopting braiding techniques to separate hair might be a good solution. For higher spatial resolution setups, labs can consider application of (or development of) suggestions as outlined in *A Guide to Hair Preparation for EEG Studies*, available online (29).

Aside from building trust with marginalized communities, hiring and training a research team with diversity in mind can bring in practitioners who can effectively relate to marginalized participants before, during, and after laboratory visits. With better familiarity of marginalized communities, researchers can identify and prevent barriers to participation, making their studies more accessible. Similarly, allocating grant money to hire a hair consultant while considering custom setups is ideal.

### 5.3 IRBs, Journals, Governing, and Foundations: Mandated Reporting

The responsibility of race and gender reporting does not simply fall on individual researchers, but also on the funding, publishing, and ethics bodies to which they are beholden. Each of these entities have a responsibility to mandate reporting of demographics and question any researchers who include race- or phenotype-based exclusion criteria in their studies. As highlighted in (7), IRBs are in place to ensure that institutional research is both rigorous and ethical. IRB personnel should receive ongoing training on the presence of racial bias in research devices and offer institutionally mandated inclusive best practices to researchers.

Similarly, funding bodies and journals should require demographic reporting and data demographic disaggregation. Foundations should invite research proposals explicitly asking the questions of the present article: who is being excluded and why, both technologically and culturally? Finally, foundations should fund innovative and equitable technologies, like the work of the team led by Sossena Wood at Carnegie Mellon University and the team led by Meryem Yücel at Boston University, both funded by Meta Reality Labs.

Pressure for change will mount with the help of concerted action-based efforts. More scientific organizations and foundations should provide support for neuroscientists and engineers via resources like the Neuroethics Framework formed by IEEE. At the Federal level, passing the Diverse and Equitable Participation in Clinical Trials (DEPICT) Act and similar legislature would help, provide the FDA with the authority to require diverse representation in clinical trials.

Though the onus of progress is collective, the authors herein embolden the entire fNIRS community to assume individual responsibility for conducting inclusive work within their own realms of influence, including as researchers, journal editors, manuscript and grant reviewers, IRB members, and leaders in their own scientific and social circles.

## ACKNOWLEDGEMENTS

JAK was supported by NIH award K00NS115331. TCP was supported by NSF GRFP #1752134. Funding for this investigation was provided by a Meta Reality Labs grant to SW and JAK.

## Note to editors

In this article, we outline the importance of demographic reporting in fNIRS articles, since there is evidence that phenotypic bias might be rampant in the field. Our work is a literature review of both neuroscience journals and biomedical optics journals to see if there is any “phenotypic exclusion” of darker skinned, darker haired, and coarse, curly haired individuals from data samples. To do this, we first look at how many articles even report demographics such as race/ethnicity and phenotype. We compare to gender reporting, a standard exemplar reporting demographic, and conclude that race, ethnicity, and phenotype are grossly underreported, making it difficult to assess if the fNIRS literature, both in neuroscience and in optics, is actually inclusive. We situate these issues with other methodological issues of fNIRS and give recommendations to researchers, funders, IRBs, and journals to mitigate these problems.

## Notes

### Competing Interest Statement

The authors have declared no competing interest.

