## Supplemental Information for "Demographic Reporting and Phenotypic Exclusion in fNIRS"

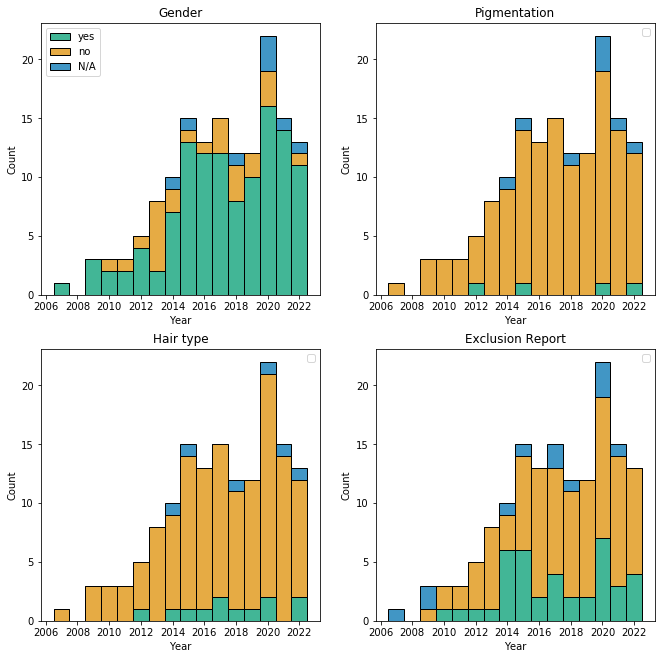

*Supplemental Figure 1: Demographic reporting over time for all 5 journals (3 biomedical optics and 2 human neuroscience). There is a marked increase of publications overall, but no apparent change in demographic reporting proportionally.*

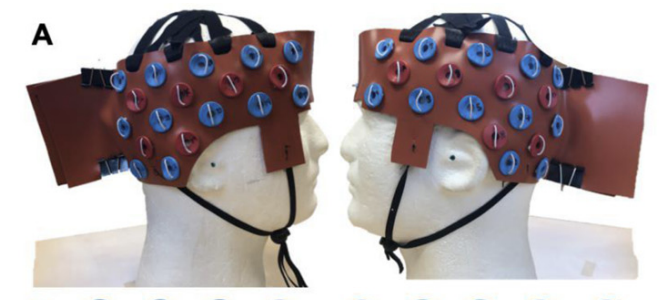

*Supplemental Figure 2. Sun et al.’s fNIRS cap configuration with custom placement embedded in a silicon strip (reproduced from Sun et al. 2022 with permission).*

*Supplemental Table 1. Full list of exclusion comments found in the 197 articles surveyed.*

| Article Type | Exclusion comment quote |
| --- | --- |
| Human Neuro journals | Three participants were excluded because fNIRS data quality was poor, mostly due to poor light shielding and excessive sunlight generating noise in the recorded light intensity signal. |
|  | Exclusion criteria for the study included: a Mini-Mental State Exam (MMSE) below 24, established cardiovascular disease at age 53, heart attack by the age of 69. |
|  | During preprocessing, the fNIRS data of two dyads (one from the high-low group and one from the low-high group) could not be viewed due to recording error. |
|  | In additional 23 infants were tested, but were excluded from the analyses due to crying or fussiness |
|  | One participant was later discovered to have had a brain tumor which was removed 2 years prior to the experimental session |
|  | 3 excluded due to noisy data across moth channels and 5 were further excluded due to exclusion of region of interest |
|  | Any displacement of the headband that compromised stability of the fit was excluded from further analysis; Intensity readings lower than a certain threshold were excluded |
|  | Two participants were excluded from all data analysis due to participant non-compliance with the study procedures. Six additional participants were excluded from fNIRS analysis due to technical issues (2 participants) or low quality fNIRS data (4 participants), leaving a final sample of 62 participants. |
|  | A participant’s entire data set was removed if > 25% of his data were ineligible and any excluded participants were replaced. |
|  | ine participants were excluded from the eye gaze analysis due to poor signal quality in the eye-tracking data |
|  | Bad fNIRS signal and technicaly issues |
|  | Technical problems and motion artifact |
|  | Noisy channels |
|  | Poor signals |
|  | Poor digitation and poor data quality |
|  | doesnt meat saturated light intensity |
|  | poor cap fit, excessive motion, and failure to complete the entire imaging protocol |
|  | exclusion criteria same as TMS (presence of seizures) |
|  | noisy data |
|  | machine malfunctioned |
|  | noise or motion artifact |
|  | data collection failure |
|  | life time history of medical illness or substance use (besides cannabis) |
|  | excluded if no coverage of bilateral prefrontal cortices |
|  | three outliers were excluded from analysis |
|  | excluded because of mild depressivity |
|  | outliers were excluded |
|  | had a lot of hair to obstruct light |
|  | noncompliance of the child technical reasons |
|  | history of mental illness |
|  | clinical reasons, alcohol/ drug dependence, traumatic brain injury, intellectual disability, "other conditions that disqualified participants as determined by investigators" |
|  | cognitive disorders, medical instability, severe aphasia, mental illness, limited range of motion in certain areas, those who received antispastic therapy |
|  | two were attributed to scheduling conflicts for the baseline assessment; one was unable to collect effective signals from fNIRS due to the participant’s thick, strong hair; one was due to having a cold during MI-BCI training; and the remaining two were unable to complete the whole study |
|  | *table presents subjects with less than 50% acceptable channels |
|  | crying or "too active" |
|  | two subjects did not pass the Cumulative sum analysis learning curve exam and were excluded from the data analysis (explained in the Supplementary Material “CUSUM scores”) |
|  | One of the healthy controls showed insufficient channel quality in the calibration stage and was, therefore, excluded from further analysis. |
|  | One subject had to be excluded due to technical problems. |
|  | Two subjects were excluded due to excessive movements during the recording. Another three subjects were excluded due to high electrode impedance in recording sessions. |
|  | Due to bad signal quality, 12 participants had to be excluded from the data analysis |
|  | We excluded subjects from the study when they presented any history of neurological or vascular disease, magnetic resonance imaging (MRI) contraindications (e.g., metal implants), or lack of a palpable inion because of neuronavigation constraints (see section “Real-Time Neuronavigation”). |
|  | 2 excluded for missing too many blocks |
|  | three had to be excluded due to significant motion artifacts and overall low signal quality. |
|  | One participant was excluded from the analysis because he reported drowsiness during the experiment. |
|  | We were unable to record fNIRS data from two participants during VFT and six participants in LMT. Therefore, these individuals were excluded from our analyses. |
|  | Two optodes, 1 and 15, were over the hairline for most participants and hence were rejected from the study. |
| Optics Journals | insufficient signal quality caused by bad scalp contacts of the optical probes and severe muscle artifacts |
|  | Since data from subject 3 have poor SNR and are severely affected by motion artifacts in half of the fNIRS data (see appendix Table 4), we could treat this subject as an outlier. Alignment accuracy without using data from subject 3 is reported in Table 1 and Fig. 8. |
|  | Contaminated data fragments caused by large head movements, unexpected behaviors, and sharp changes in fNIRS signals were removed by a visual inspection [9, 30]. Five children in the ASD group and two children in the TD group with too short retention time (i.e., less than 8 min) or with one or more poor signal-to-noise ratio channels were excluded from further data analysis. |
|  | Consequently, 13 participants were invited in the experiment; however, data from two participants were excluded from further processing due to incorrect operation during the fNIRS experiment. |
|  | Three of them were later excluded from further analysis due to poor probe contact or signal quality, resulting in a final number of 6-6 subjects in the two groups (3-3 female). |
|  | One participant’s data was discarded because of large motion artifacts. Another was discarded because of bad optical coupling between the optodes and the scalp. The third was rejected because of poor data quality resulting from the subject’s relatively thick, black hair. |
|  | Bad channels (for example due to loose contact to skin) were excluded from further analysis. |
|  | Data from two participants were excluded from further processing due to participant loss and data-recording failure, respectively. |
|  | Subjects with a history of neurological trauma or psychiatric disorders were excluded |
|  | Exclusion criteria were any psychiatric or neurological disorder or current medication |
|  | Participants were excluded from the imaging analysis if they did not complete all the necessary tasks or did not fit this study’s inclusion criteria |
|  | 4 participants were excluded as they did not understand our instructions for the experiment. |
|  | excluded channels not people because of indicated poor or inadequate optical signals that usually resulted from hair absorption and/or imperfect contact of optodes to the scalp |
|  | Three of the subjects were excluded due to poor optical contact between the head probe and scalp on at least one measurement channel |
|  | Potential participants were excluded if they had a history of seizure, paroxysm, or hormonal, metabolic, circulatory, psychiatric or neurological illness, or were currently pregnant. |
|  | One of the participants did not complete the study |
|  | In order to remove the the motion artifact from our study and make sure the data were qualified for analysis and interpretation, we introduced the same training procedures as our previous studies on children with ASD. Before the experiment, we randomly sampled 24 children with ASD. After training, 3 children with ASD did not meet the standard of experiment and 1 child with ASD still moved his head to follow the red dot during the experiment |
|  | One control subject was excluded from analysis due to poor contact between the source detectors and the scalp |
|  | two infants were tested but refused the fNIRS head probe. No infants refused the ECG sensors. |
|  | only three participants among them were able to finish the experiments to determine the task duration; Because the experiments lasted for a long time, an exhausted participant could not participate in some experiments; The highest classification accuracy was obtained using PCA for participants 1, 3, 6, 7, and 8, because the LDA could not be applied to the high-dimensional feature vector because of the singular scatter matrix problem |
|  | gross systemic noises/artifacts in the signal or unexpected death during the experiment |
|  | Therefore, Ch. 7 is considered as the most active channel in MC. If the mean value from some subjects (Ch. 7) is less than or equal to zero, the subject is supposed to be excluded from further analysis: In the MC experiment, four subjects (3, 4, 16, and 17) were excluded. |
|  | data from one child (7 years old, the youngest recruited in this study) were discarded due to large motion-induced artifacts |
|  | subject did not return due to flashing lights |
|  | three participants had consistently noisy fNIRS data across all the channels in motor stimulation sessions. It was presumably due to dense and/or dark-colored hair, which could lead to loose contact of fNIRS optodes on the head or high light absorption44 if some hair was left between the optode and scalp. Thus these three participants were excluded from data analysis |
|  | ability for repeat testing, handedness |
|  | five subjects were excluded from any further analysis because they either showed large head movements, or the data were too noisy due to hair obstruction |
